# Unsupervised Whole-Genome Representation Learning Captures Bacterial Phenotypes

**DOI:** 10.1101/2025.04.01.646674

**Authors:** Cameron Dufault, Alan M. Moses

## Abstract

Shifting from hand-crafted to learned representations of data has revolutionized fields like natural language processing and computer vision. Despite this, current approaches to bacterial phenotype prediction from the genome rely on training machine learning models on hand-crafted features, often binary indicators or counts of the presence of different conserved genomic elements and protein domains. Defining these shared elements and domains as our “genomic element vocabulary”, we tokenize entire bacterial genomes as sequences of these conserved elements and take advantage of advances in long-context language modeling to perform self-supervised whole-genome representation learning (WGRL). Through multi-task pretraining on a phylogenetically diverse dataset of hundreds of thousands of bacterial genomes, we present a genomic language model which produces representations of input genomes with features predictive of a broad range of phenotypes. We assess the quality of the learned representations through k-nearest neighbours prediction of 25 bacterial phenotypes, finding our WGRL representations more predictive than standard protein domain presence/absence representations for 23/25 different phenotypes. We additionally find the WGRL representations are robust to both poor genome assembly quality and incompleteness. Through learning the relationships between evolutionarily conserved genomic elements with self-supervised long-context language modeling, we demonstrate the first approach for extracting general-purpose whole-genome representations while preserving gene order.

## 1 Background AND Introduction

At its most fundamental level, information in the genome is encoded through just 4 nucleotides. However, like letters in an alphabet, sequences of these nucleotides within the genome can be classified into various genomic elements. These include genes (both coding and non-coding), regulatory regions, and repetitive elements. Interactions between these genomic elements and their products occurs at all levels of information flow in the central dogma, forming complex networks like metabolic pathways and gene regulatory networks essential for building and maintaining life.

This complexity has frustrated expectations that sequencing of whole genomes would lead to understanding of how genomes encode an organism’s phenotype. The advent of metagenomic sequencing has made performing microbial genotype-to-phenotype prediction crucial. Most newly-discovered microbial species cannot be cultured (Feldbauer et al. (2015)) and are known only from their genome sequences, and estimates of the number of prokaryotic species are as high as 1 trillion (Locey & Lennon (2016)). However, knowledge-based approaches to predicting phenotypes using mechanistic understandings of cellular processes are limited to well-studied species and phenotypes (Karlsen et al. (2023)). This has led to the development of data-driven supervised machine learning approaches to performing prokaryotic genotype-to-phenotype prediction.

To create training examples, these approaches require expressing whole prokaryotic genomes as numerical representations. Currently, genomes are most commonly represented as vectors with thousands of hand-crafted features indicating the presence/absence or copy number counts of evolutionarily conserved genomic elements like clusters of orthologous genes (COGs, Tatusov et al. (1997)) or protein domain families, generally Pfam domains (Mistry et al. (2021); Karlsen et al. (2023)). This high dimensionality means that when few labeled examples are available supervised models may learn to make predictions by exploiting phylogenetic correlations rather than understanding the mechanisms behind a given phenotype (Li et al. (2023)).

As shifting from manually engineered to learned representations of data has led to breakthroughs in the abilities of machine learning models across many domains, representation learning of genomic sequences has been an area of intense research interest. However, there have been very few efforts towards learning the representations of *whole genome* sequences used for genotype-to-phenotype prediction. Many existing approaches to genomic sequence representation learning make use of natural language processing (NLP) techniques to create genomic language models (gLMs) trained using self-supervised learning (SSL). Typically, gLMs are trained to model interactions within DNA sequences tokenized at the level of single nucleotides or short k-mers, but this approach cannot scale to whole genomes. Recent long-context gLMs (Nguyen et al. (2024)) have been trained to model sequences of up to 131 kilobases in length at single-nucleotide resolution, but this remains orders of magnitude smaller than the average bacterial genome (3.65 Mb, diCenzo & Finan (2017)).

Some gLMs model interactions between genomic elements rather than nucleotides themselves, capturing longer-range interactions. These higher-order gLMs treat whole genes or protein domains as tokens, utilizing the fact that prokaryotic genomes are dense with coding genes belonging to the families of evolutionarily conserved genomic elements whose presence is often used as features for genotype-to-phenotype prediction. Miller et al. (2022) and Tan et al. (2024) utilize word2vec (Mikolov et al. (2013)) to learn contextualized representations of COGs and protein domains from millions of annotated microbial genome contigs, however a window size of just 5 tokens was used. Rather than learning token embeddings from scratch, Hwang et al. (2024) embed each coding gene within a dataset of millions of short microbial contigs (15-30 genes long) using an existing protein language model (ESM-2, Lin et al. (2023)), stack these embeddings, then train a BERT-based language model (Devlin et al. (2018)) to fill in masked genes based on their context within each contig. Using a similar approach but with much longer context, Li et al. (2024) demonstrate, to the best of our knowledge, the only approach for directly modeling interactions across whole genomes while preserving order. They also stack gene embeddings from ESM-2, but create whole-genome inputs by concatenating all contigs in a genome together. Using this approach, a BERT-based model was trained on the task of predicting habitat specificity on a dataset of 29,089 whole bacterial genomes. While Li et al. (2024) do not demonstrate producing generalizable representations, they demonstrate an order conserving, gene-based approach to learning representations of whole genomes for phenotype prediction. Importantly, this accounts for how bacterial genomes are organized. While human genomes are only *∼* 1.5% coding genes, on average 87% of bacterial genomes are (Land et al. (2015)). Additionally, gene order in prokaryotic genomes has been found to have functional importance, with clusters of genes beyond operons being maintained over evolution (Tamames (2001)). The position along a chromosome where a bacterial gene is found as well as its copy number has also been shown to influence its expression (Bryant et al. (2014); Dryselius et al. (2008)).

Here, we extend this idea further to demonstrate unsupervised whole-genome representation learning (WGRL) as a method for producing whole-genome representations. Leveraging the increase in availability of high-quality assembled bacterial genomes as well as advances in NLP, we use self-supervised learning to train a long-context gLM on a large and diverse collection of assembled and annotated whole bacterial genomes, tokenized according to a vocabulary of conserved genomic elements common across the tree of life. We validate our WGRL approach produces representations of bacterial genomes which are more predictive of phenotypes than standard Pfam domain/presence absence representations through k-NN prediction of 25 different phenotypes. We additionally assess whether our learned representations are robust to genome incompleteness and assembly quality.

## 2 Methods

### 2.1 Dataset creation

A large and diverse repository of consistently annotated, high-quality, and non-redundant whole bacterial genome assemblies for WGRL training were obtained through RefSeq (O’Leary et al. (2016)). Almost all bacterial genomes in RefSeq are annotated using NCBI’s prokaryotic genome annotation pipeline (PGAP, Haft et al. (2024)) which identifies coding sequences, pseudogenes, CRIPSR repeats, and non-coding RNA, including tRNA, rRNAs, and small ncRNAs. rRNAs and small ncRNAs are additionally searched with RFAM (Kalvari et al. (2021)) models. We downloaded annotation features (GFF) files, and protein (FASTA) files for each of the 371175 (as of August 12 2024) bacterial genome assemblies in RefSeq. HMMER3 (Eddy (2011)) was then used to scan each of the *∼* 280 million unique protein sequences for Pfam domains (Pfam-A, v35.0). Only domain hits with domain score greater than 10 were assigned to each protein and stored in a dictionary.

We express every bacterial genome assembly downloaded from RefSeq as a sequence of conserved genomic elements and protein domains. For each assembly, the GFF file was read line-by-line, with genomic elements being added to a growing genomic element sequence in the order they appear. We annotated each coding gene with their Pfam domains, as they can be assigned to a significantly higher percentage of proteins in the bacterial proteome than COGs (Lobb et al. (2020)). For each coding sequence, Pfam domains assigned to the protein are looked-up in the domain hits dictionary. The names of the domains are then added to the genomic elements sequence in the order they appear in the protein. If domains overlap, only the domain with the higher domain score is added. For each non-coding gene encountered, the identifier of the RFAM family assigned to the gene is added, except in the case of tRNA where the specific product is added (e.g., “tRNA-Met”). A “pseudogene” token is also added for each pseudogene annotation, as well as tokens for each annotated repeat belonging to a named family (e.g., “CRISPR”). Finally, punctuation tokens are added to reflect the structure of the genome. A “contig_start” token is added when each new contig in the assembly is encountered. “protein_start” and “protein_end” tokens are added for each coding gene, surrounding any Pfam domains found. To reflect the double-stranded nature of the genome, “+” and “–” tokens are also added before every token that is found on the opposite strand of the previous token.

After conversion to genomic element sequence format, genomes were split into training, test, and validation datasets according to phylogeny to allow for assessing the ability of the trained model to generalize to genomes of different phylogenetic distances to the training set. To increase diversity, the training dataset was filtered to reduce highly similar sequences, then sequences from underrepresented species were upsampled. Details of this procedure can be found in A.3.

*∼* 23000 unique tokens were found in the training dataset. To create a genomic element vocabulary containing only tokens found across the tree of bacterial life, we selected only tokens appearing in genomes from at least 1000 different genera, out of the 3459 genera represented. This gave a vocabulary of 3562 tokens, of which 3515 were Pfam domains, 35 were non-coding RNA (both tRNA products and Rfam families), 5 were our previously defined “punctuation” tokens, 5 were special tokens, and the remaining two indicated pseudogenes and CRISPR repeats. Each genomic element sequence in the dataset was tokenized according to this vocabulary, with unknown words mapped to the [UNK] token and [CLS] and [SEP] tokens being added to the ends of each sequence.

### 2.2 Model Architecture and Training Procedure

Figure 1 provides an overview of the gLM architecture and training scheme. The model is based on BiMamba (Schiff et al. (2024)), which extends Mamba (Gu & Dao (2023)) to support bi-directional sequence modeling by processing sequences in both the forward and reverse direction, generating two projections of the same sequence which are then added together. A bi-directional, Mamba-based architecture was chosen as genomic elements have both upstream and downstream interactions, and the tokenized sequences were too long to apply full attention (median of 15498 tokens across all bacterial RefSeq genomes). Data augmentations were applied to reduce potential overfitting and encourage the model to learn representations robust to genome incompleteness and poor assembly quality. We randomly cut contigs longer than 1000 tokens such that the two new contigs are at least 100 tokens in length. The number of times long contigs in each sequence were cut was randomly chosen between 1 and 5. However, contigs shorter than 1000 tokens were never cut, meaning if after a round of the cuts there were no contigs in the sequence longer than 1000 tokens remaining, no more cuts were made. After cutting and shuffling contigs (as they have no known order), we randomly truncated 85% of genomes, taking a random window of 40% to 60% of the overall sequence.

**Figure 1:**
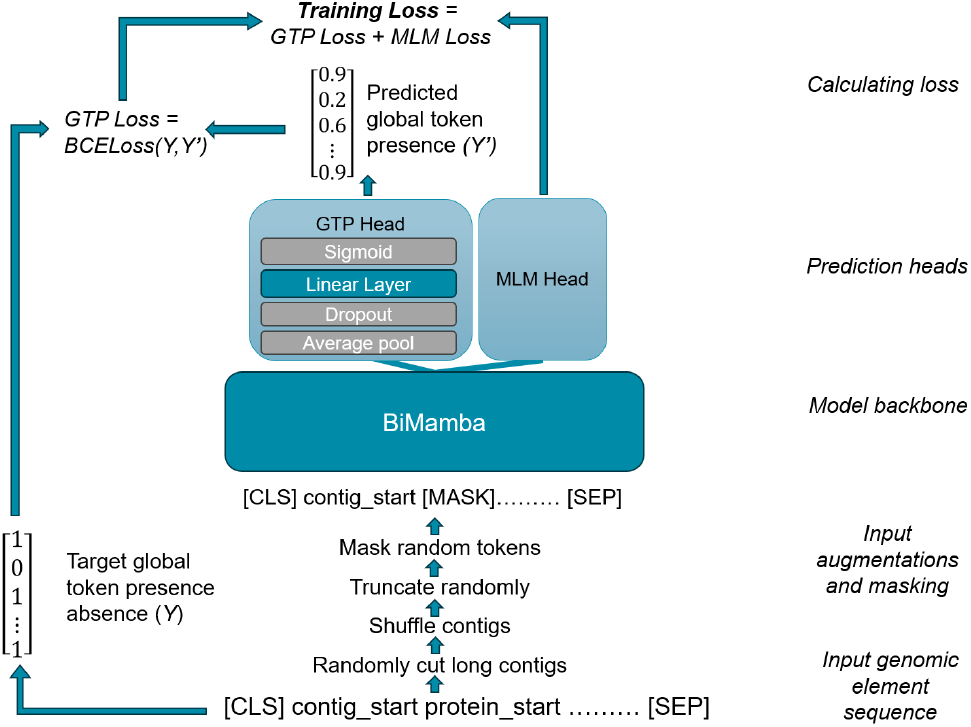
Overview of gLM architecture and training procedure.

A novel multi-task pretraining objective was used. In addition to masked-language modeling (MLM, using the standard BERT masking strategy from Devlin et al. (2018)), we introduced a global token presence (GTP) task which requires the model to predict which tokens were present in the genome sequence before both random masking and random sequence truncation were applied. The addition of the GTP task was motivated by noticing the standard bag-of-words Pfam presence/absence genome representations were surprisingly predictive of many phenotypes. The GTP task head makes its predictions using the final hidden layer representation of the model, averaged across the sequence to be a *d*_*hidden_size*_ vector. This head consists of a dropout layer followed by linear classification layer and a sigmoid activation layer, producing a *d*_*vocab_size*_ vector of token probabilities. The output of the GTP head is then compared to the target binary vector indicating the originally present tokens, with the binary cross entropy loss between the predicted and target vectors calculated. This GTP task loss is then added with the MLM loss. We trained two variations of the gLM for performing WGRL, one with combined MLM and GTP task loss, and one which only learned from the MLM loss. Further details on model parameters and training can be found in A.2.

### 2.3 K-nearest neighbours BacDive phenotype prediction

#### 2.3.1 Downloading data and choosing phenotypes to predict

To assess whether the wgrl representations were predictive of bacterial phenotypes, 15926 genomes with a diverse array of phenotype labels were sourced from bacdive (schober et al. (2024)). as previously described, proteins in each genome were annotated with pfam domains and each genome expressed as a genomic element sequence. 20 binary and 5 continuous phenotype labels were able to be associated with at least 1000 downloaded bacdive genomes. these phenotypes related to strain morphology (motility, gram stain, cell shape, multicellular complex forming, cell length, and cell width), growth conditions (optimal temperature, optimal ph, optimal salinity), physiology and metabolism (sporulation, oxygen tolerance, acetoin production, nitrate reduction, glucose assimilation, urea hydrolysis, methyl-red test, indole test, voges-proskauer test, and 0129 resistance test) and environment (soil, animal, and aquatic). both continuous and binary labels were associated with cell length and width, and oxygen tolerance was divided into two binary labels, is anaerobe and is aerobe. a detailed list of phenotypes can be found in a.1 tables 3 and 4.

### 2.3.2 performing k-nearest neighbours prediction

K-nearest neighbours (k-NN) classification is a widespread approach to evaluating the quality of representations learned via self-supervised learning (Lee et al. (2023)), with weighted k-NN being a common variation used (Caron et al. (2021); Wu et al. (2018)) where the classification of a query point is more influenced by closer neighbours than those farther away. To evaluate the quality of our learned representations, we used a weighted k-NN classifier to predict the 20 binary bacterial phenotypes and a weighted k-NN regressor to predict the 5 continuous phenotypes, and compared the performance against the standard Pfam domain presence/absence representations.

Learned WGRL representations for all 15926 BacDive genomes were extracted from the final layer of the trained gLMs and averaged across the sequence to be *d*_*hidden size*_ vectors. We used the set of tokens found in each genomic element sequence to create Pfam domain presence/absence representations, with a binary indicator for each of the 20894 tokens appearing in at least 20 different genomes in the RefSeq training dataset, 20793 of which were Pfam domains. With both our learned and standard representations for each genome, the genomes were split into training and test datasets 5 times using 5-fold cross validation, such that each genome was in the test dataset one time. Over each fold, we assessed the performance of weighted k-NN for predicting each of the 25 phenotypes using the whole-genome representations. If more than one genome from the same species had a phenotype label, only one randomly chosen genome from that species was retained. Cosine distance was used as the distance metric, with neighbours in the training dataset being weighted by the inverse of their cosine distance to the query point. As there are many potential choices of k, we calculated performance on all phenotypes with k values of 1, 5, 10, 15, 20, 25, and 30.

To assess whether the WGRL representations are robust to poor assembly quality, we compared the k-NN phenotype prediction performance between well and poorly assembled genomes in the Bac-Dive dataset. Genomes stored by NCBI are assigned one of four assembly levels based on the quality of the genome assembly, with “contig” level assemblies being the worst. Contig level genomes in the BacDive dataset have a median of 45 contigs, while across all others the median is 16. We selected all “contig” level assemblies as our poorly assembled genomes and all other genomes as well assembled, and separately tracked the performance on the poorly assembled and well assembled genomes. To assess whether the WGRL representations are robust to incompleteness, we also evaluated the k-NN phenotype prediction performance when test set genomes were reduced to 50%, 33%, 25%, and 12.5% of their original length before their representations created.

## 3 Results

### 3.1 WGRL representations are more predictive than standard approach

The results of k-NN BacDive phenotype prediction evaluation show that the WGRL representations extracted from the gLM are more predictive of a broad range of phenotypes than the standard Pfam presence/absence representations. Taking the average performance on all tasks, WGRL representations beat the standard representations at all k values except k=1, the worst performing value of k. This is seen for both binary classification and regression tasks (Figure 2a). Taking a closer look at the performance on each task when k=15 (chosen because it is the midpoint between k=10 and k=20, the values resulting in the highest average performance for each representation) we can see that this performance improvement is the result of a consistent edge by the WGRL representations over the standard representations across most tasks (Figure 2b). Looking at MCC (Table 1) and *r*_*s*_ performance (Table 2) for each task shows the WGRL representations are more predictive than the standard representation approach for 23/25 phenotypes when k=15. Importantly, our learned representations demonstrated high predictive performance for phenotypes with both high and low phylogenetic signal (the degree to which the phenotype is shared with phylogenetically similar species). A previous study found that pH preference and salinity preference, two phenotypes our model outperformed the standard representation on, do not show significant phylogenetic signal Barberán et al. (2017).

**Table 1:** MCC performance of k-NN phenotype prediction for each binary phenotype (k=15). Error is the standard error of the mean MCC across 5-fold cross validation.

| Phenotype | WGRL MCC (k=15) | Pfam presence/absence MCC (k=15) |
| --- | --- | --- |
| acetoin production | 0.330±0.016 | **0.368±0.002** |
| binary length | **0.601±0.031** | 0.523±0.033 |
| binary width | **0.444±0.028** | 0.436±0.026 |
| complex forming | **0.736±0.023** | 0.711±0.033 |
| glucose assimilation | **0.485±0.012** | 0.454±0.027 |
| Gram stain | 0.955±0.002 | **0.956±0.001** |
| indole test | **0.527±0.019** | 0.424±0.017 |
| is aerobe | **0.808±0.007** | 0.797±0.007 |
| is anaerobe | **0.925±0.005** | 0.907±0.007 |
| is animal | **0.938±0.004** | 0.913±0.008 |
| is aquatic | **0.923±0.007** | 0.914±0.004 |
| is soil | **0.906±0.018** | 0.878±0.018 |
| methyl-red test | **0.467±0.027** | 0.403±0.041 |
| motility | **0.675±0.004** | 0.659±0.007 |
| nitrate reduction | **0.448±0.010** | 0.430±0.011 |
| resistance to O/129 | **0.550±0.032** | 0.521±0.029 |
| rod or coccus shaped | **0.733±0.014** | 0.689±0.021 |
| sporulation | **0.893±0.008** | 0.885±0.013 |
| urea hydrolysis | **0.441±0.016** | 0.430±0.021 |
| Voges-Proskauer test | **0.384±0.009** | 0.381±0.020 |

**Table 2:** Spearman’s rank correlation coefficient (*r*_*s*_) of k-NN phenotype prediction for continuous phenotypes (k=15). Error is the standard error of the mean *r*_*s*_ across 5-fold cross validation.

| Phenotype | WGRL $r_s$ (k=15) | Pfam presence/absence $r_s$ (k=15) |
| --- | --- | --- |
| average length | **0.518±0.019** | 0.501±0.016 |
| average width | **0.473±0.017** | 0.458±0.013 |
| minimum optimal temperature | **0.534±0.008** | 0.516±0.018 |
| optimal pH | **0.474±0.017** | 0.455±0.017 |
| optimal salinity concentration | **0.630±0.008** | 0.605±0.011 |

**Table 3:**
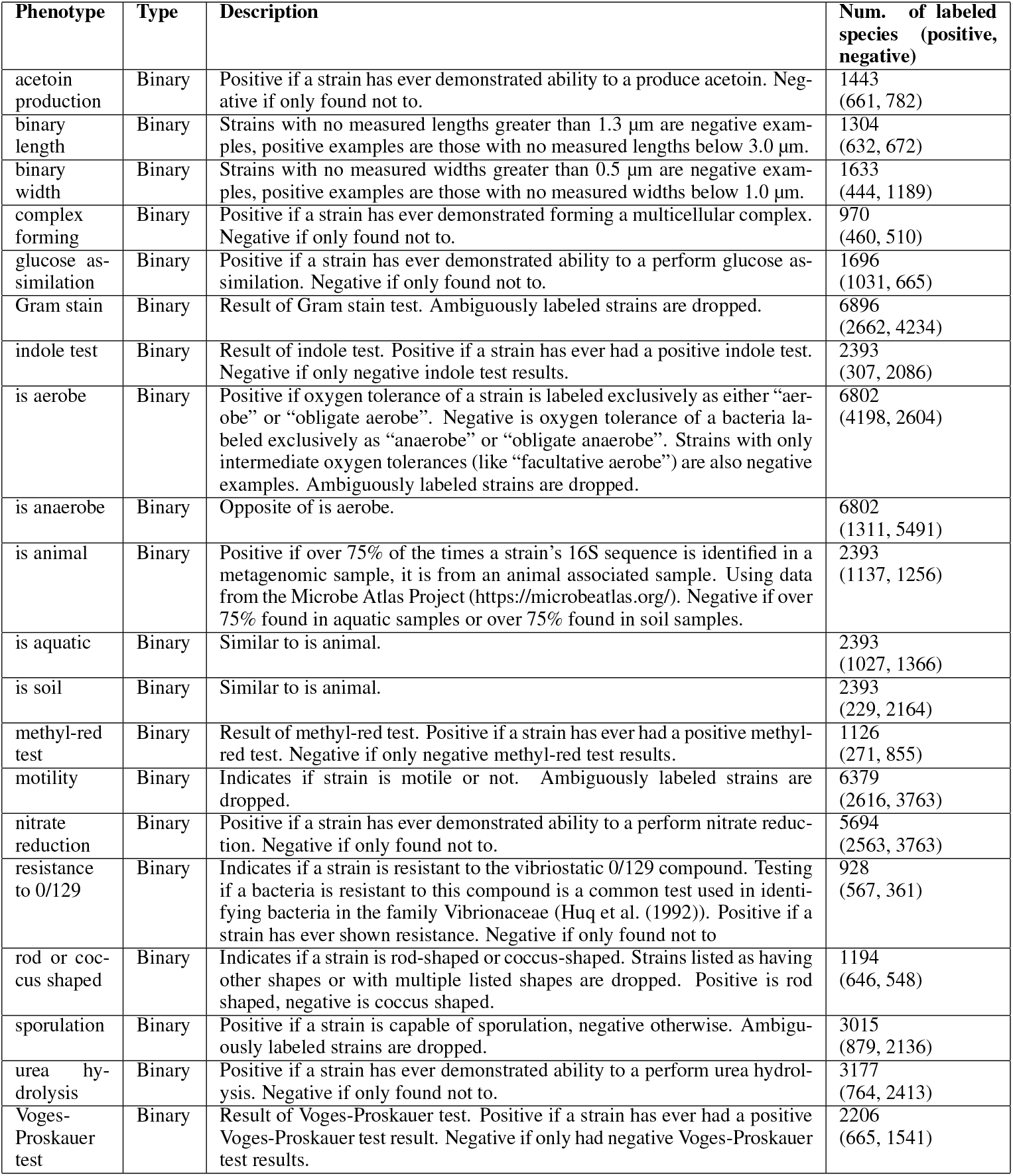
List of predicted binary phenotypes.

**Table 4:** List of predicted continuous phenotypes.

| Phenotype | Type | Description | Num. of labeled species (positive, negative) |
| --- | --- | --- | --- |
| average length | Continuous | The average measured cell length. If BacDive lists cell length as a range, the center of this range is chosen. If BacDive has multiple cell length values, they are averaged. | 3938 |
| average width | Continuous | The average measured cell width. If BacDive lists cell width as a range, the center of this range is chosen. If BacDive has multiple cell width values, they are averaged. | 3828 |
| minimum optimal temperature | Continuous | The minimum measured optimal growth temperature ($^{\circ}\text{C}$) of a strain. If BacDive lists optimal growth temperature as a range, the minimum of this range is chosen. If more than one listed optimal growth temperature, we take smallest. | 3652 |
| optimal salinity concentration | Continuous | The measured optimal salinity (NaCl %) for cell growth. If BacDive lists optimal salinity as a range, the center of this range is chosen. If more than one listed optimal salinity value, the first is chosen. | 2575 |
| optimal pH | Continuous | The measured optimal pH for cell growth. If BacDive lists optimal pH as a range, the center of this range is chosen. If more than one listed optimal pH value, the first is chosen. | 3654 |

**Figure 2:**
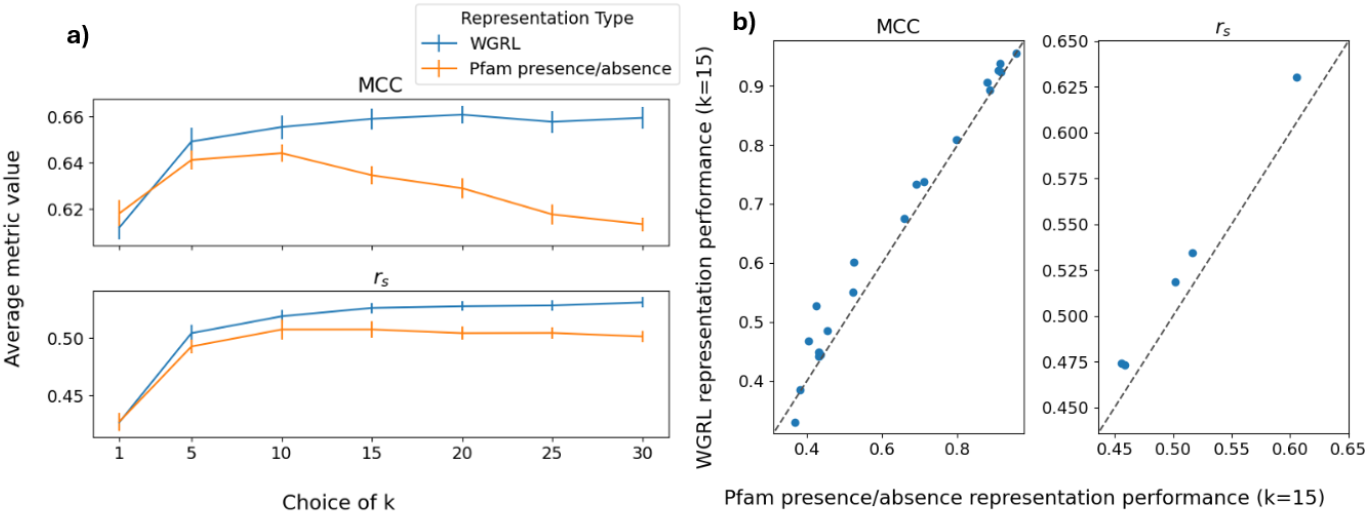
Comparison of WGRL and Pfam presence/absence representation performance for k-NN phenotype prediction. a) Average performance across all phenotype predictions for each representation type at different values of k. b) Comparison of the performance of the two representations when k=15. The dotted lines represent equal performance.

### 3.2 WGRL representations are robust to assembly quality and incompleteness

The performance of the wgrl representations was robust to poor genome assembly quality. classification performance differed only slightly, with mcc performance on the 20 binary tasks when using well assembled genomes being on average only 0.00940 higher than when only using poorly assembled genomes. performance on 8 tasks increased while it dropped for 12 others (figure 3).

**Figure 3:**
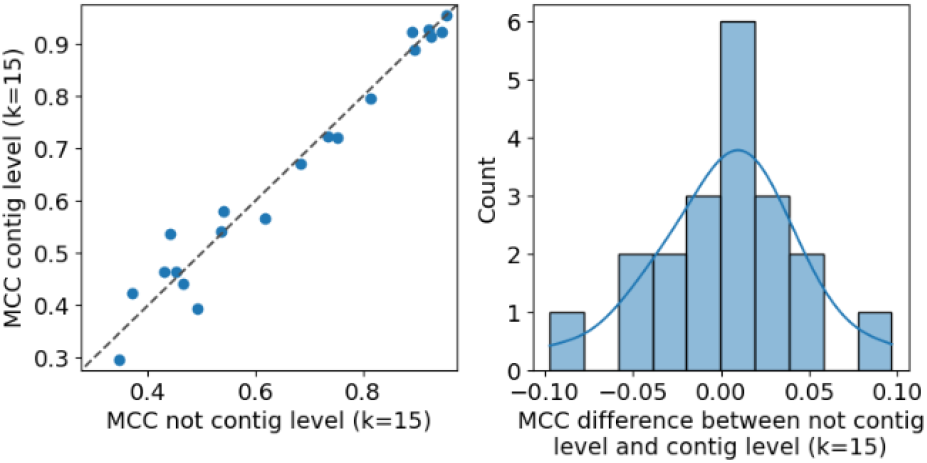
Comparison of MCC performance on well assembled versus poorly assembled genomes. the left plot shows the performance on each task with each set of genomes (well-assembled on the x-axis and poorly assembled on the y-axis). the right plot shows the distribution of the shift in MCC (well assembled minus poorly assembled)

Figure 4 shows the effect on k-nn prediction performance of genome incompleteness. as expected, performance using both learned and standard representations increases as a higher fraction of the full genome is used. we find that the wgrl representations outperform the standard representations at all levels of genome completeness, with the difference in performance between the two being most pronounced when genomes are 25%, 33%, and 50% complete. this indicates high-quality wgrl representations can be produced even from highly incomplete genomes.

**Figure 4:**
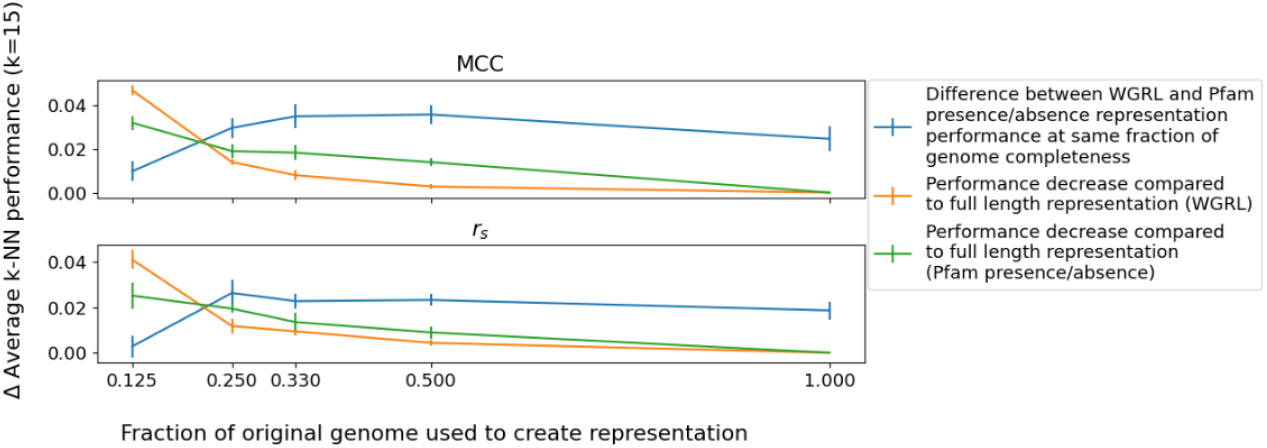
Comparison between WGRL representations and Pfam presence/absence representations for k-NN (k=15) phenotype prediction with genomes of different fractions of completeness.

### 3.3 global token presence task drives phenotype predictive representations

As shown in Figure 5, the average k-NN performance on the phenotype prediction tasks using the gLM trained with GTP + MLM loss is higher at all choices of k than the gLM trained identically except only using the MLM loss. Representations from the MLM only gLM perform similarly to the standard Pfam presence/absence representations, although the highest average levels of performance occur at higher k choices than the standard Pfam representations. A comparison with the performance of representations from a randomly initialized BiMamba model gives an idea of how much the gLMs learned from pretraining through self-supervised WGRL.

**Figure 5:**
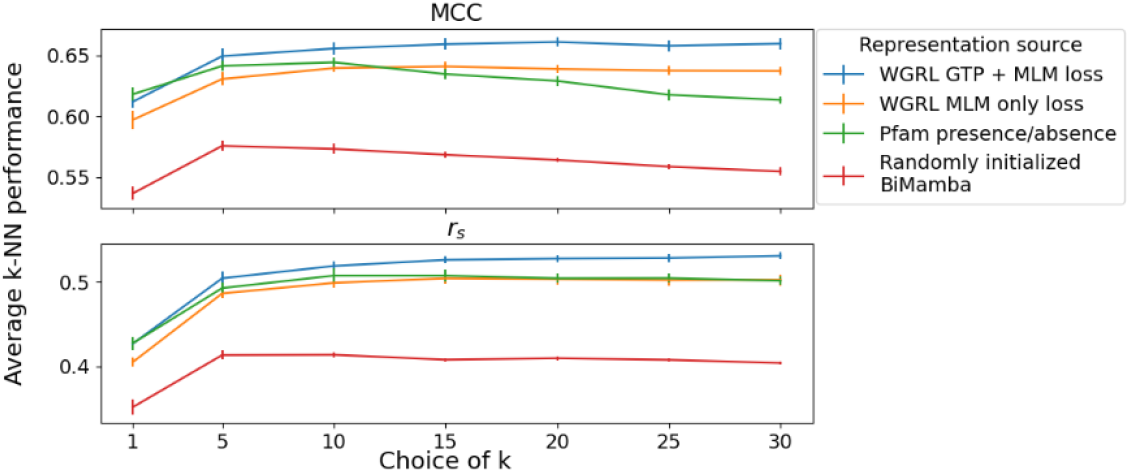
Average phenotype prediction performance across all binary classification and regression tasks for genome representations from different sources.

## 4 Discussion

### 4.1 Summary

We introduced unsupervised whole-genome representation learning, an approach for learning from unlabeled genome sequence data which produces whole genome representations predictive of bacterial phenotypes. After tokenizing a diverse dataset of hundreds of thousands of bacterial genomes as sequences of a vocabulary of genomic elements, we used self-supervised learning to train a long-context gLM to model the relationships between these evolutionarily conserved and functionally distinct units spread across bacterial genomes. After training, our model produced representations of bacterial genomes more predictive of 23/25 phenotypes than standard Pfam domain presence/absence representations, and which are robust to poor genome quality and incompleteness.

As it has in other problem domains, using learned representations rather than manually engineered features may lead to dramatically improved bacterial genotype-to-phenotype prediction. There is a vast and constantly growing amount of publicly available bacterial genome sequence data. This data can be leveraged through self-supervised learning to produce representations which capture the biological meaning of the complex network of interactions in the genome. Our method, which we believe is the first method for learning general-purpose whole-genome representations while preserving gene order, demonstrated the ability to learn from this data to create informative representations. Our results present a potential path for improved bacterial phenotype prediction, an increasingly important task as the amount of unculturable bacterial species known from their genomes alone continues to increase.

### 4.2 Limitations and future work

Whole-genome representations from gLMs using genomic element level tokens are not suitable for predicting phenotypes caused by nucleotide level variations, such as certain antibiotic resistance phenotypes (Davis et al. (2016)). By not modeling at the nucleotide level, widespread genomic features like GC content and codon bias, both of which are known to be correlated with some phenotypes, are also lost (Barnum et al. (2024)). Calculating these features separately and appending them to the learned representations may be a way of incorporating DNA level information when making phenotype predictions with WGRL representations.

While we focus on proteins and non-coding genes, there are other genomic elements in bacterial genomes like operons, promoter sequences, and transcription factor binding sites which play important roles in the regulation of bacterial gene expression. Future work could involve using existing tools for annotating bacterial genomes which can identify promoters, transcription factor binding sites, different types of mobile genetic elements, and biosynthetic gene clusters (Jung et al. (2024)) to increase the diversity of tokens available to learn from. Additionally, a median of 239 coding sequences in every bacterial genome in RefSeq could not be associated with any Pfam domains, and thus no information about these proteins was encoded in the genomic element sequence. Alternative approaches to representing these proteins should be explored in future work.

We did not systematically compare the performance of our learned representations against existing state-of-the-art genotype-to-phenotype approaches. We have not found benchmark datasets which new bacterial genotype-to-phenotype prediction methods can readily be tested on, and as such making direct comparisons between the performance of different existing methods is difficult. Even for predicting the same phenotype, most existing approaches have widely varying data sources, dataset sizes, and control for phylogeny in different ways. For this reason, we limited ourselves to assessing the quality of our learned representations against a standard Pfam presence/absence domain representation using a k-NN approach, as this required no parameter tuning. Future evaluations should also compare against other standard whole-genome representation approaches like COGs presence/absence and k-mer counts. We hope the development of deep learning models for bacterial phenotype prediction will drive the creation of fair benchmarks for model comparison.

### Meaningfulness Statement

A meaningful representation of life is one which can distill relevant biological concepts from poorly structured and/or noisy data. Ideally, learning of meaningful representations is guided by knowledge of biological information organization. Our work contributes to increasing meaning in genome representation learning by 1) modeling interactions between functionally distinct and evolutionary conserved units of information in the genome and 2) representing the entire genome, the biologically complete genotype. We show that this approach leads to improved phenotype prediction, one of the major challenges in biology.

## Data and Code Availability

Code and instructions for downloading data, training models, and performing evaluation available at: https://github.com/dufaultc/bacterial-whole-genome-representation

## A Appendix

### A.1 Phenotype details

#### A.2 Model details

Each gLM used to produce learned representations had 16 layers, a hidden size of 768, and 69,532,906 parameters. The models were each trained for 4 epochs, batch size of 32, and a maximum learning rate of 4.0 *×* 10^*−*4^. We used the AdamW optimizer with 0.1 weight decay and *β*_1_ and *β*_2_ values of 0.9 and 0.95, as well as a cosine learning rate scheduler with a 10% warmup ratio. Training each model took 8 days on an NVIDIA RTX A6000 GPU.

#### A.3 Dataset splitting, filtering, and upsamping

To create the test set, 5% of the 3983 different genera represented in the corpus were selected and genomes belonging to these genera moved to the test set, followed by genomes belonging to 5% of all remaining species and then 5% of all remaining assemblies. To create the validation dataset, 5% of the remaining genera were moved to the validation dataset. The remaining genomes (n=265285) were used to build the training dataset. As highly studied species were overrepresented, the training dataset was filtered to maximize the level of diversity. Genomes which were not type-strains and which had high average nucleotide identity (above 99.5%) or high coverage (above 95%) by a typestrain were dropped. Next the dataset was subset by selecting as many genomes as possible of each species, to a maximum of 100000 total genomes. Genomes from species with less than 18 examples were then up-sampled according to their level of representation, to a maximum of 10 times for species with just one example. This resulted in the final training dataset having 535967 genomes.

